## Supplementary Information for "A Commander-independent function of COMMD3 in endosomal trafficking"

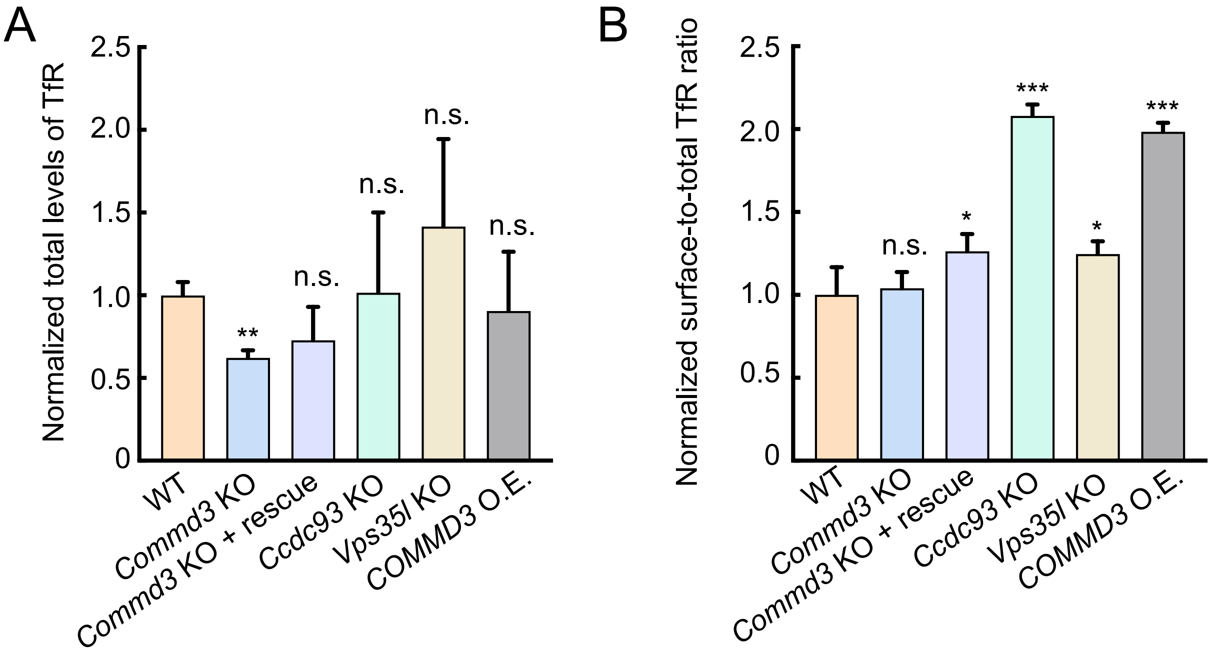


**Figure S1. Quantification of transferrin receptor (TfR) in wild-type (WT) and mutant cell lines. (A)** Quantification of total TfR levels in the indicated preadipocyte cell lines based on protein intensities from immunoblots, analyzed using ImageJ. Data were normalized to WT cells. Data are presented as mean ± SD from three biological replicates. ** $P < 0.01$; n.s., $P > 0.05$ (one-way ANOVA). **(B)** Normalized surface-to-total TfR ratio. Surface TfR levels, measured by flow cytometry, were normalized to total protein expression. Data are presented as mean ± SD from three biological replicates. * $P < 0.05$; *** $P < 0.001$; n.s., $P > 0.05$ (one-way ANOVA). Related to Figures 4 and 5.

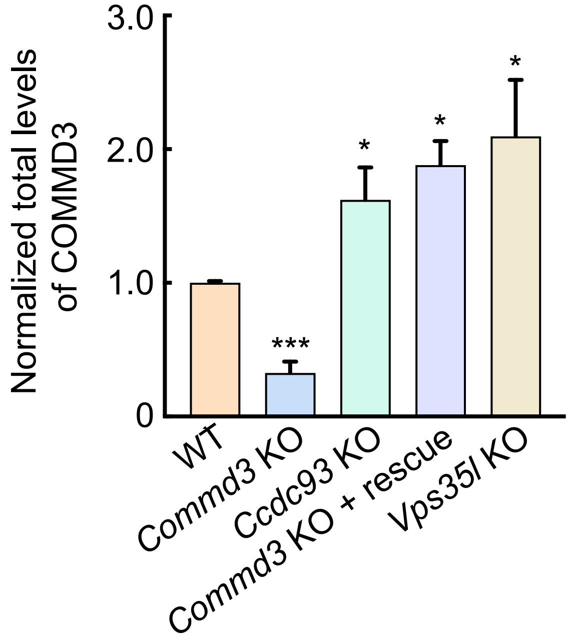


**Figure S2. Quantification of total COMMD3 levels**. Total COMMD3 levels in the indicated preadipocyte cell lines were quantified based on protein intensities from immunoblots, analyzed using ImageJ. Data were normalized to WT cells. Data are presented as mean ± SD from three biological replicates. *** $P < 0.001$; * $P < 0.05$ (one-way ANOVA). Related to Figure 5.

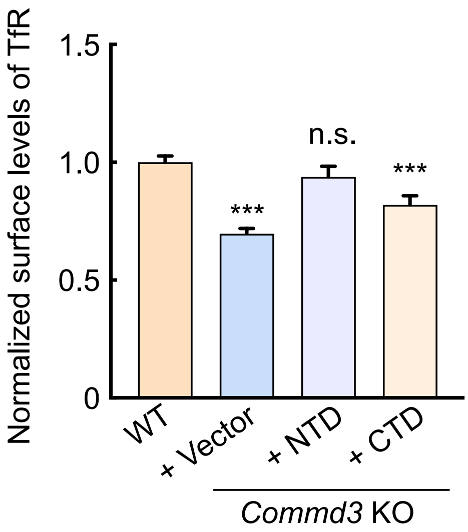


**Figure S3. The CTD of COMMD3 is unable to rescue TfR surface levels.** Normalized surface levels of TfR were measured by flow cytometry in the indicated preadipocyte cell lines. Data were normalized to WT cells. Results are presented as mean ± SD from three biological replicates. *** $P < 0.001$; n.s., $P > 0.05$ (one-way ANOVA). Related to Figure 6.
